# Falco: A quick and flexible single-cell RNA-seq processing framework on the cloud

**DOI:** 10.1101/064006

**Authors:** Andrian Yang, Michael Troup, Peijie Lin, Joshua W. K. Ho

## Abstract

**Summary:** Single-cell RNA-seq (scRNA-seq) is increasingly used in a range of biomedical studies. Nonetheless, current RNA-seq analysis tools are not specifically designed to efficiently process scRNA-seq data due to their limited scalability. Here we introduce Falco, a cloud-based framework to enable paralellisation of existing RNA-seq processing pipelines using big data technologies of Apache Hadoop and Apache Spark for performing massively parallel analysis of large scale transcriptomic data. Using two public scRNA-seq data sets and two popular RNA-seq alignment/feature quantification pipelines, we show that the same processing pipeline runs 2.6 – 145.4 times faster using Falco than running on a highly optimised single node analysis. Falco also allows user to the utilise low-cost spot instances of Amazon Web Services (AWS), providing a 65% reduction in cost of analysis.

**Availability:** Falco is available via a GNU General Public License at https://github.com/VCCRI/Falco/

**Supplementary information:** Supplementary data are available at *BioRXiv* online.

## 1 Introduction

Major advancements in single-cell technology have resulted in an increasing interest in single-cell level studies, particularly in the field of transcriptomics [12]. Single-cell RNA sequencing (scRNA-seq) offers the promise of understanding transcriptional heterogeneity of individual cells, allowing for a clearer understanding of biological process [14, 8, 4, 11].

Each scRNA-seq experiment typically generates profiles of hundreds of cells, which is a magnitude larger than the typical amount of data generated by standard bulk RNA-seq experiments. Current RNA-seq processing pipelines are not specifically designed to handle such a large number of profiles. To fully realise the potential of scRNA-seq, we need a scalable and efficient computational solution. The premise of our solution is that state-of-the-art cloud computing technology, which is known for its scalability, elasticity and pay-as-you-go payment model, can allow for a highly efficient and cost-effective scRNA-seq analysis.

There are a number of existing cloud-based next-generation sequencing bioinformatics tools based on the Hadoop framework, an open source implementation of MapReduce [5], or the Spark framework [17]. Halvade, written in Hadoop MapReduce, is designed to perform variant calling of genomic data from FASTQ files, though it also offers support for transcriptomic analysis [6]. SparkSeq [16] and SparkBWA [1], both written in Spark, offers interactive sequencing analysis of BAM files and alignment of FASTQ files respectively. These tools have limitations in the context of scRNA-seq analysis. Of the three tools, only SparkSeq allows for multi-sample analysis, although SparkSeq itself is also limited as it does not perform alignment, which is the main bottleneck in sequence analysis.

Here we use a different approach to utilising cloud-based big data technology. Our framework – Falco – is a framework that allows users to ‘plug-in’ their chosen RNA-seq alignment, quality control, preprocessing and feature quantification tools, and enable the resulting pipeline to run multi-sample analysis of large-scale transcriptomic data on the cloud. Falco utilises Amazon Elastic MapReduce (EMR), a big data processing service for deploying managed Hadoop and Spark clusters on the Amazon Web Services (AWS) cloud.

## 2 Framework

The Falco framework consists of a splitting step, an optional pre-processing step and the main analysis step. The first step, the splitting step, is a MapReduce job which splits FASTQ input files stored in the Amazon S3 storage service into multiple smaller FASTQ files. In the case of paired-end reads, the two reads are combined into a single record to ensure that paired-end reads are processed together. The splitting process is performed in order to increase the level of parallelism in analysis and normalise the performance of tools as each chunk will have the same maximum uncompressed size of 256 MB.

The next step in the pipeline is an optional step for performing pre-processing of reads, such as adapter trimming and filtering reads based on quality. The pre-processing step is another MapReduce job which performs pre-processing of the split FASTQ files using any pre-processing tools chosen by the user. The user is asked to supply a shell script with commands to run their selected pre-processing tools, that is then called by the MapReduce job.

The final step of the pipeline is the main analysis step. It performs alignment and quantification of reads using the Spark framework. It was designed such that any RNA-seq alignment and quantification tools can be used within the Falco framework. In the current implementation, each split FASTQ file can be aligned using either STAR [7] or HISAT2 [9] and quantified using either featureCounts [13] or HTSeq [2]. By default, STAR and featureCount will be used for alignment and quantification, however the framework accepts any combination of the tools. The returned gene counts per split are then reduced (i.e., merged) to obtain the total read counts per gene in each sample. The gene count matrix is produced and stored into Amazon S3 storage. Aside from the gene counts, the analysis step also returns selected mapping and quantification reports generated by the selected alignment and quantification tools as well as optional RNA-seq alignment metrics from Picard tools [3].

As part of the pipeline, a script is provided to simplify the creation of the EMR cluster and configure the required software and references on the cluster. Similarly, each of the steps also has a corresponding submission script which will upload the files required for the step and submit the step to the EMR cluster for execution.

### 2.1 Customising Falco framework

The Falco framework allows the user to add custom alignment and/or quantification tools beyond what is provided by default. Instructions are provided in the github wiki which will take the user through the steps required to add their selected tool(s) to the framework. It is expected that the user has moderate to advanced Python proficiency in order to perform customisation of the framework.

To ensure that the output of Falco matches that of non-Falco execution, the tools must be compatible with divide-and-conquer approach. Examples of tools which are not compatible with Falco approach include TopHat2 [10] and StringTie [15] as those tools uses information from the entire read for performing calling and quantification, respectively. The divide-and-conquer approach used by Falco means that the tools only have partial information from the entire read and thus the output will not necessarily be the same.

## 3 Evaluation

To evaluate the performance of Falco, the runtime of two popular RNA-seq pipelines, STAR followed by featureCounts (S+F), and HISAT2 followed by HTSeq (H+H), is evaluated using two scRNA-seq data sets with and without using the Falco framework. A number of realistic scenarios for analysis in a single computing node were devised – from the naïve single processing approach to a highly parallelised approach. Furthermore, to demonstrate the scalability of Falco, EMR clusters with increasing numbers of core nodes (from 10 to 40) were used to show the effect of adding more computational resources on the runtime of Falco.

In all the comparison, the AWS EC2 instance type used for computation (core node for EMR) is r3.8xlarge (32 cores, 244GB of RAM and two 320GB SSDs). For Falco’s EMR cluster, a single r3.4xlarge (16 cores, 122GB RAM) was used as the master node for scheduling jobs and managing the cluster. The EMR cluster uses Amazon EMR release 4.6, which contains Apache Hadoop 2.7.2 and Apache Spark 1.6.1, and takes 16 minutes for initialisation and configuration in all cluster configurations used.

Two recently published scRNA-seq datasets were used for evaluation. The first dataset (SRA accession: ERP005988), is a mouse embryonic stem cell (mESC) single cell data containing 869 samples of 200 bp paired-end reads, totalling to 1.28 × 10^12^ sequenced bases, stored in 1.02 Tb of gzipped FASTQ les [11]. The second dataset (SRA accession: SRP057196), is a smaller human brain single cell data containing 466 samples of 100 bp paired-end reads, totalling to 2.95 × 10^11^ sequenced bases and 213.66 Gb of gzipped FASTQ files [4].

Comparing the performance of a single node, with different parallelisation approaches, against Falco shows that running the S+F pipeline on Falco results in a speedup of 2.6x (10 nodes vs 16 processes) to 33.4x (40 nodes vs 1 process) for the mouse dataset and 5.1x (10 nodes vs 16 processed) to 145.4x (40 node vs 1 process) for the human dataset. For the H+H pipeline, Falco gives a speedup of 2.5x (10 nodes vs 16 processes) to 58.4x (40 nodes vs 1 process) and 4.0x (10 nodes vs 16 processes) to 132.5x (40 nodes vs 1 process) for the mouse and brain datasets respectively (Table 1). The disparity in the speed-up between the two datasets is due to different pre-processing tools being employed, with the human dataset utilising more pre-processing steps in the original publication [4]. We also note that the gene expression quantification produced by a given pipeline is the same regardless of whether the Falco framework was used.

**Table 1:** scRNA-seq processing time with or without Falco

| System | Nodes | Mouse - embryonic<br>stem cell (hours) |  | Human - brain<br>(hours) |  |
| --- | --- | --- | --- | --- | --- |
|  |  | S+F* | H+H* | S+F* | H+H* |
| Standalone | 1 (1 process) | 93.7 | 233.6 | 154.7 | 198.8 |
|  | 1 (12 processes) | 21.1 | 32.6 | 16.4 | 19.6 |
|  | 1 (16 processes) | 18.5 | 28.4 | 13.6 | 16.2 |
| Falco | 10 | 7 | 11.2 | 2.7 | 4.1 |
|  | 20 | 4.1 | 6.4 | 1.6 | 2.3 |
|  | 30 | 3.3 | 4.8 | 1.4 | 2.0 |
|  | 40 | 2.8 | 4.0 | 1.1 | 1.5 |

\*S+F = STAR for aligner and featureCounts for quantification; H+H = HISAT2 for aligner and HTSeq for quantification. Standalone number of processes indicates the number of FASTQ file pairs that are processed in parallel. Timing for Falco includes initialisation and configuration time which are approximately 16 minutes.

For the scalability comparison, it can be seen that the runtime of the pipeline decreases with increasing cluster size (Table 1), though the trend is gradual rather than linear. Analysis of the runtime for each step in the framework shows a similar gradual decrease in runtime for pre-processing and analysis steps (Supplementary Figure 2). For the splitting step, a different trend is seen where there is little to no decrease in runtime for cluster size ≥ 20 nodes. The lack of speed up for splitting is due to the number of executors exceeding the number of files to be split and the limitation of time taken to split large files as the distribution of file size in both test datasets is uneven (Supplementary Figure 1).

To save cost, EMR allows for the usage of reduced price *spot* computing resources. The spot prices fluctuate depending on the availability of the unused computing resource and the spot instance is obtained by supplying a bid for the resource. The use of spot instances for analysis provide a substantial saving of around 65% compared to using on-demand instances (Table 2 and 3). The trade-off with using spot instances is that the computing resource could be terminated should the market price for that resource exceed the user’s bid price.

**Table 2:** Falco cost analysis: on-demand vs. spot instances for STAR + featureCounts

| Dataset | Cluster size | Time<br>(hours) | On-demand<br>cost (USD) | Spot cost<br>(USD) | % Savings |
| --- | --- | --- | --- | --- | --- |
| Mouse - | 10 node | 8 | 247.20 | 85.67 | 65.34 |
| embryonic | 20 node | 5 | 301.00 | 99.09 | 67.08 |
| stem cell | 30 node | 4 | 258.00 | 115.71 | 55.15 |
|  | 40 node | 3 | 356.40 | 114.11 | 67.98 |
| Human - | 10 node | 3 | 92.70 | 32.13 | 65.34 |
| brain | 20 node | 2 | 120.40 | 39.64 | 67.08 |
|  | 30 node | 2 | 179.00 | 57.86 | 67.68 |
|  | 40 node | 2 | 237.60 | 76.08 | 67.98 |

Time rounded up to whole hour including cluster startup. Price used for r3.8xlarge instance is USD\$2.660/hr (on-demand price) and USD\$0.64/hr(average spot price for June 2016).

**Table 3:** Falco cost analysis: on-demand vs. spot instances for HISAT2 + HTSeq

| Dataset | Cluster size | Time<br>(hours) | On-demand<br>cost (USD) | Spot cost<br>(USD) | % Savings |
| --- | --- | --- | --- | --- | --- |
| Mouse - | 10 | 13 | 401.70 | 139.10 | 65.37 |
| embryonic | 20 | 7 | 421.40 | 138.60 | 67.11 |
| stem cell | 30 | 5 | 447.50 | 144.50 | 67.71 |
|  | 40 | 4 | 475.20 | 152.00 | 68.01 |
| Human - | 10 | 5 | 154.50 | 53.50 | 65.37 |
| brain | 20 | 3 | 180.60 | 59.40 | 67.11 |
|  | 30 | 2 | 179.00 | 57.80 | 67.71 |
|  | 40 | 2 | 237.60 | 76.00 | 68.01 |

Time rounded up to whole hour including cluster startup. Price used for r3.8xlarge instance is USD\$2.660/hr (on-demand price) and USD\$0.64/hr(average spot price for June 2016).

## 4 Summary

Falco is a cloud-based framework that enables massively parallelised sequence alignment, quality control, and feature quantification of single-cell transcriptomic data in AWS cloud-computing environment.

